# Transcription activation domains of the yeast factors Met4 and Ino2: tandem activation domains with properties similar to the yeast Gcn4 activator

**DOI:** 10.1101/228536

**Authors:** Derek Pacheco, Linda warfield, Michelle Brajcich, Hannah Robbins, Jie Luo, jeff Ranish, Steven Hahn

**Affiliations:** Fred Hutchinson Cancer Research Center, Seattle WA 98109; University of Washington, School of Medicine, Seattle, WA 98195; The institute for Systems Biology, Seattle WA 98109

## Abstract

Eukaryotic transcription activation domains (ADs) are intrinsically disordered polypeptides that typically interact with coactivator complexes, leading to stimulation of transcription initiation, elongation and chromatin modifications. Here we examine the properties of two strong and conserved yeast ADs: Met4 and Ino2. Both factors have tandem ADs that were identified by conserved sequence and functional studies. While AD function from both factors depends on hydrophobic residues, Ino2 further requires key conserved acidic and polar residues for optimal function. Binding studies show that the ADs bind multiple Med15 activator binding domains (ABDs) with a similar order of micromolar affinity, and similar but distinct thermodynamic properties. Protein crosslinking shows that no unique complex is formed upon Met4-Med15 binding. Rather, we observed heterogeneous AD-ABD contacts with nearly every possible AD-ABD combination. Many of these properties are similar to those observed with the yeast activator Gcn4, which forms a large heterogeneous, dynamic, and fuzzy complex with Med15. We suggest that this molecular behavior is common among eukaryotic activators.

## Introduction

Transcription activators play essential roles in gene regulation and regulation of activator function is often the endpoint of many signaling pathways, serving to modulate transcription in response to developmental pathways, growth, stress, and other environmental signals (1, 2). The targeting of multiple activators in different combinations to gene regulatory regions leads to diverse patterns of gene regulation. Activators can enhance RNA Polymerase II transcription through binding to coactivator complexes such as Mediator, SAGA, TFIID, Swi/Snf and NuA4, complexes that contact the basal transcription machinery and/or function to modify chromatin (3, 4). Most eukaryotic activators contain separate DNA binding and transcription activation domains (ADs) (3, 5). Unlike DNA binding domains, which are usually structurally ordered, eukaryotic ADs are intrinsically disordered, lacking a stable structure (6-10).

Many types of intrinsically disordered proteins bind their targets via short linear motifs, 3-10 residue sequences that function as recognition sites for enzymes such as kinases, acetylases or methylases or as substrates for peptide binding domains such as SH2, SH3 and 14-3-3 domains (11-14). In contrast, different ADs have little primary sequence similarity, although they are often enriched for acidic, proline and glutamine residues (15, 16). At least part of this sequence bias is due to overrepresentation of these residues in intrinsically disordered proteins (17). Known ADs vary in length from a ~5 residue sequence motif to nearly 100 residues (18-20). Mutations created within ADs have shown that their function can be remarkably resistant to mutagenesis, although hydrophobic and sometimes acidic residues are critical for activity (3,21).

One of the best characterized activators is yeast Gcn4, a transcription factor that activates a large set of genes in response to metabolic stress (22), (23). Gcn4 contains tandem acidic ADs of unrelated sequence and interacts with the coactivators Mediator, SAGA, NuA4, TFIID, and Swi/Snf (6, 18, 19, 24-30). Binding of Gcn4 to the Mediator tail module subunit Med15 occurs via multiple heterogeneous interactions between the tandem ADs and up to 4 activator-binding domains (ABDs) on Med15 termed KIX, ABD1, ABD2 and ABD3 (27, 28). The measured individual binding interactions are dynamic with half-lives on the low millisecond timescale (6). Combined biochemical and structural analysis showed that the interaction between Gcn4 ADs and Med15 is “fuzzy” as Gcn4 binds to the Med15 activator-binding domains in multiple orientations (6, 18) and the fuzzy nature of this complex is conserved upon interaction of the tandem ADs with full length Med15. This binding mechanism can explain how many activators bind multiple unrelated targets using a variety of AD sequences. In contrast, several well-characterized activators are known from structural studies to bind their targets using a different mechanism that utilizes a higher affinity and more specific protein-protein interface (7, 8, 31, 32).

To explore whether other activators have properties similar to Gcn4, we used molecular, genetic, and biochemical approaches to characterize two strong yeast activators, Met4 and Ino2. Both factors have tandem acidic ADs that are moderately conserved in closely related yeasts but have primary sequences that are unrelated to each other and to Gcn4. Despite these sequence differences, Met4, Ino2, and Gcn4 have similar function in transcription activation assays, require Med15 for activation of Mediator Tail dependent promoters, and both ADs bind Med15 activator-binding domains with low micromolar affinities. These and other results suggest that Gcn4, Met4 and Ino2 use a similar strategy to bind Mediator that involves a large, dynamic and fuzzy protein interface.

## Methods

### Strains and Plasmids

All yeast strains and primary plasmids used in this work are listed in **Table S1**.

### Cell growth assays and measurement of steady state mRNA levels

Yeast strains were grown in duplicate to an OD_600_ of 0.5–0.8 in 2% (wt/vol) dextrose synthetic complete Ile-Val-Leu medium at 30 °C. Cells were induced with 0.5 μg/mL SM for 90 min to induce amino acid starvation (27), RNA was extracted and assayed in triplicate by RT-quantitative PCR, and the results were analyzed as described (27).

### Quantitation of in Vivo AD-Gcn4 Levels

Cells (1.5 mL) from the cultures used for the above mRNA analysis were pelleted and incubated in 0.1M NaOH for 5 minutes at room temp. Cells were then pelleted and resuspended in 1× lithium dodecyl sulfate sample buffer (Life Technologies) containing 50 mM DTT and treated and analyzed as previously described (18).

### Protein purification

All proteins were expressed in BL21 (DE3) RIL *E. coli.* Med15 constructs were expressed and purified as described in Tuttle et al. (20). Ino2 1-41-(GS)_3_-96-160 ((GS)_3_ is the linker: GSGSGS) and Met4 72-160 constructs were expressed as N-terminal His6-SUMO-tagged proteins (Invitrogen). Cells were lysed in 50 mM HEPES pH 7.0, 500 mM NaCl, 40 mM Imidazole, 10% glycerol, 1 mM PMSF, 5 mM DTT and purified using Ni-Sepharose High Performance resin (GE Healthcare). Proteins were eluted in 50 mM HEPES pH 7.0, 500 mM NaCl, 500 mM Imidazole, 10% glycerol, 1 mM PMSF, 1 mM DTT. Purified SUMO-tagged proteins were concentrated using 10K MWCO centrifugal filters (Millipore), diluted 10x in 50 mM HEPES pH 7.0, 500 mM NaCl, 40 mM Imidazole, 10% glycerol, 1 mM PMSF, 5 mM DTT, and digested with SUMO protease for 3-5 hrs at room temperature using ~1:800 protease:protein ratio. Cleaved His6-Sumo tag was removed using Ni-Sepharose. Peptides were further purified by chromatography on Source 15Q (GE Healthcare) using a 50-350 mM NaCl gradient. To remove residual SUMO tag in the sample due to co-elution on Source 15Q, Ino2 peptides were purified over SUMO-1(CR) resin (Nectagen) and collected in the flow through. All proteins were further purified using size exclusion chromatography on Superdex 75 10/30 (GE Healthcare). Proteins used in fluorescence polarization and isothermal titration calorimetry were eluted in 20 mM KH_2_PO_4_, pH 7.5, 200 mM KCl. Proteins used in crosslinking-Mass spectrometry (CL-MS) were eluted in PBS pH7.2. The concentration of the purified proteins was determined by UV/Vis spectroscopy with extinction coefficients calculated with ProtParam {Gasteiger:2005hs}.

### FP and ITC binding experiments

Ino2 1-41-(GS)_3_-96-160 and Met4 72-160 used in fluorescence polarization were labeled with Oregon Green 488 dye (Invitrogen) as described in (27). FP measurements were conducted using a Beacon 2000 instrument as described in (27), with concentrations of Med15 spanning 0-200 μM (ABD3) or 0-125 μM (ABD123, KIX + ABD123). FP data was analyzed using Prism 7 (Graphpad Software, Inc.) to perform non-linear regression analysis using the one-site total binding model Y=Bmax*X/(Kd+X) + NS*X + Background where Y equals arbitrary polarization units and X equals Med15 concentration.

ITC titrations were performed using a Microcal ITC200 Microcalorimeter in 20 mM KH_2_PO_4_, pH 7.5, 200 mM KCl as described in (6). The following protein concentrations were used: Med15 6-90 (79.7 μM) vs. Ino2 1-41-(GS)_3_-96-160 (1.32 mM); Med15 6-90 (79.7 μM) vs. Met4 72-160 (1.27 mM); Med15 158-238 (111 μM) vs. Ino2 1-41-(GS)_3_-96-160 (2.59 mM); Med15 158-238 (117 μM) vs. Met4 72-160 (1.27 mM); Med15 277-368 (113 μM) vs. Ino2 1-41-(GS)_3_-96-160 (1.32 mM); Med15 277-368 (59.7 μM) vs Met4 72-160 (732 μM); Med15 484-651 (111 μM) vs.

Ino2 1-41-(GS)_3_-96-160 (1.32 mM); Med15 484-651 (119 μM) vs Met4 72-160 (1.12 mM). The following parameters were the same for all runs: cell temperature 22°C, reference power 11 μcal/sec, initial delay 120 sec, stir speed 1000 rpm, injection spacing 180 sec, filter period 5 sec, and injection rate 0.5 μl/sec. Activator was added over 16 injections (injection 1 = 0.4 μl, injections 2-16 – 2.55 μl). Calorimetric data were plotted and fit with a single binding site model using Origin 7.0 software (Microcal).

### EDC crosslinking and MS sample preparation

50 μg of Med15 1-651 Δ239-272, Δ373-483 (KIX + ABD123) was mixed with 3x molar excess of Ino2 1-41-(GS)_3_-96-160 or Met4 72-160. Samples were incubated with 15 mM (Met4) or 10 mM (Ino2) EDC and 2 mM Sulfo-NHS (Thermo Scientific) in 50 μl total volume PBS pH7.2 (Met4) or 150 μl total volume PBS pH 6.5 for 2 hours at room temperature. Proteins were processed for MS analysis similarly to described in Tuttle et al (20). Protein samples were reduced with 50 mM TCEP and denatured with 8 M urea at 37°C for 15 min. The samples were then alkylated in the dark at 37°C with 15 mM iodoacetamide for 1 hour. The samples were then diluted 10-fold with 100 mM ammonium bicarbonate and digested with Glu-C (20:1 w/w) over night at 37°C. Samples were then digested with trypsin (1:15 w/w) overnight at 37°C. Digested samples were purified by C18 chromatography (Waters), eluted in 80% acetonitrile 0.15 trifluoroacetic acid, and dried in a speedvac.

### MS and data analysis

EDC–cross-linked peptides were analyzed on a Thermo Scientific Orbitrap Elite at the Proteomics facility at the Fred Hutchinson Cancer Research Center and data were analyzed as described in (33). Spectra were manually evaluated using the COMET/Lorikeet Spectrum Viewer (Trans-Proteomic Pipeline) as described in (33).

## Results

### Mediator tail-dependence of transcription activators

As a first step in exploring the mechanism of yeast ADs in comparison to Gcn4, we tested the activity and coactivator dependence of several previously characterized transcription factors. Segments from 7 transcription factors with published AD function were fused to the N-terminus of the Gcn4 central region linker + Gcn4 DNA binding domain (Gcn4 residues 124-281) and tested for activation of two Gcn4-dependent promoters: *ARG3* and *HIS4* (both TATA-containing – defined as TATAWAW (34)). The expression of these AD-Gcn4 derivatives was from low copy ARS, CEN-containing plasmids under control of the Gcn4 regulatory region with ^~^1 kb DNA upstream from the Gcn4 ORF containing all known Gcn4 transcription and translational regulatory regions. The regions of these factors tested for function were: Met4 residues 1-160 (35); Ino2 residues 1-160 (36); Pdr1 residues 901-1068 (37); Hap4 residues 321-490 (38); Gal4 residues 840-881 (39, 40); Rtg3 residues 1-250 and 375-486 (41). Fusion proteins contained a C-terminal triple Flag epitope tag to monitor protein expression (18). Gcn4 synthesis and activity is induced in response to amino acid starvation, so activity of these chimeric activators was measured 90 min after addition of sulfometuronmethyl (SM), an inhibitor of Ile and Val biosynthesis, to the cell growth media (27).

**Figure 1.**
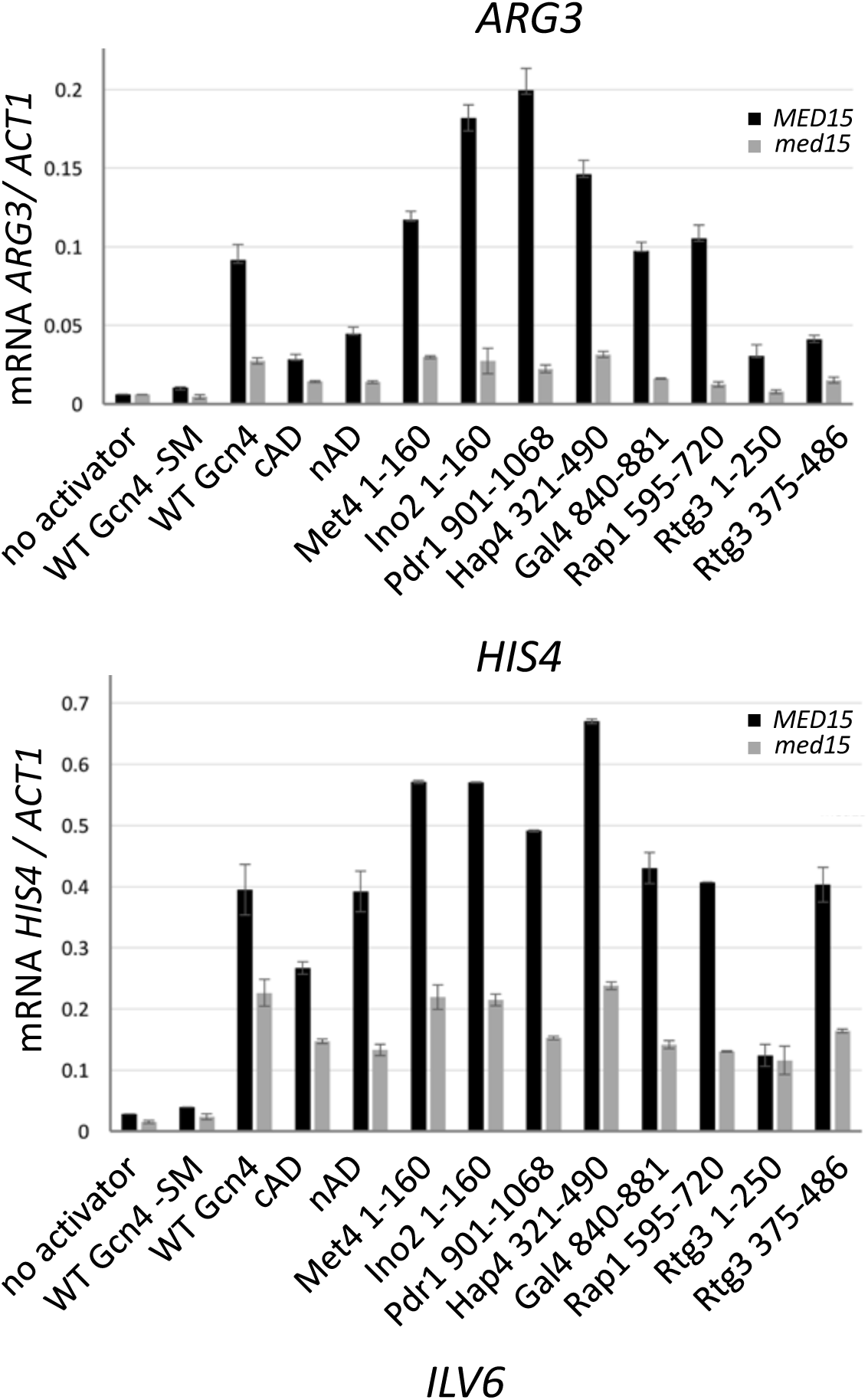
Activity of yeast transcription factor AD – Gcn4 DBD fusions at two Gcn4 inducible genes. Previously defined AD regions were fused to Gcn4 residues 125-281 and expressed under control of the Gcn4 gene regulatory region. Gcn4 inducing conditions were initiated by addition of SM for 90 min. and mRNA levels from *ARG3* and *HIS4* were quantitated by RT qPCR. Measurements were made in both *MED15* and *med15Δ* strains as indicated.

Figure 1 shows that, when fused to the Gcn4 DBD, all these ADs function to activate transcription at *ARG3* and *HIS4*, although their relative activity depends on the specific promoter. Met4, Ino2, Pdr1 and Hap4 are strong ADs at both promoters, comparable or better than wild type Gcn4. The two Rtg3 ADs have different relative activity, depending on the promoter, with the C-terminal AD having the most activity at *HIS4*. Western analysis showed that all proteins were expressed and that the level of expression did not correlate with AD function (**Fig 2A**).

**Figure 2.**
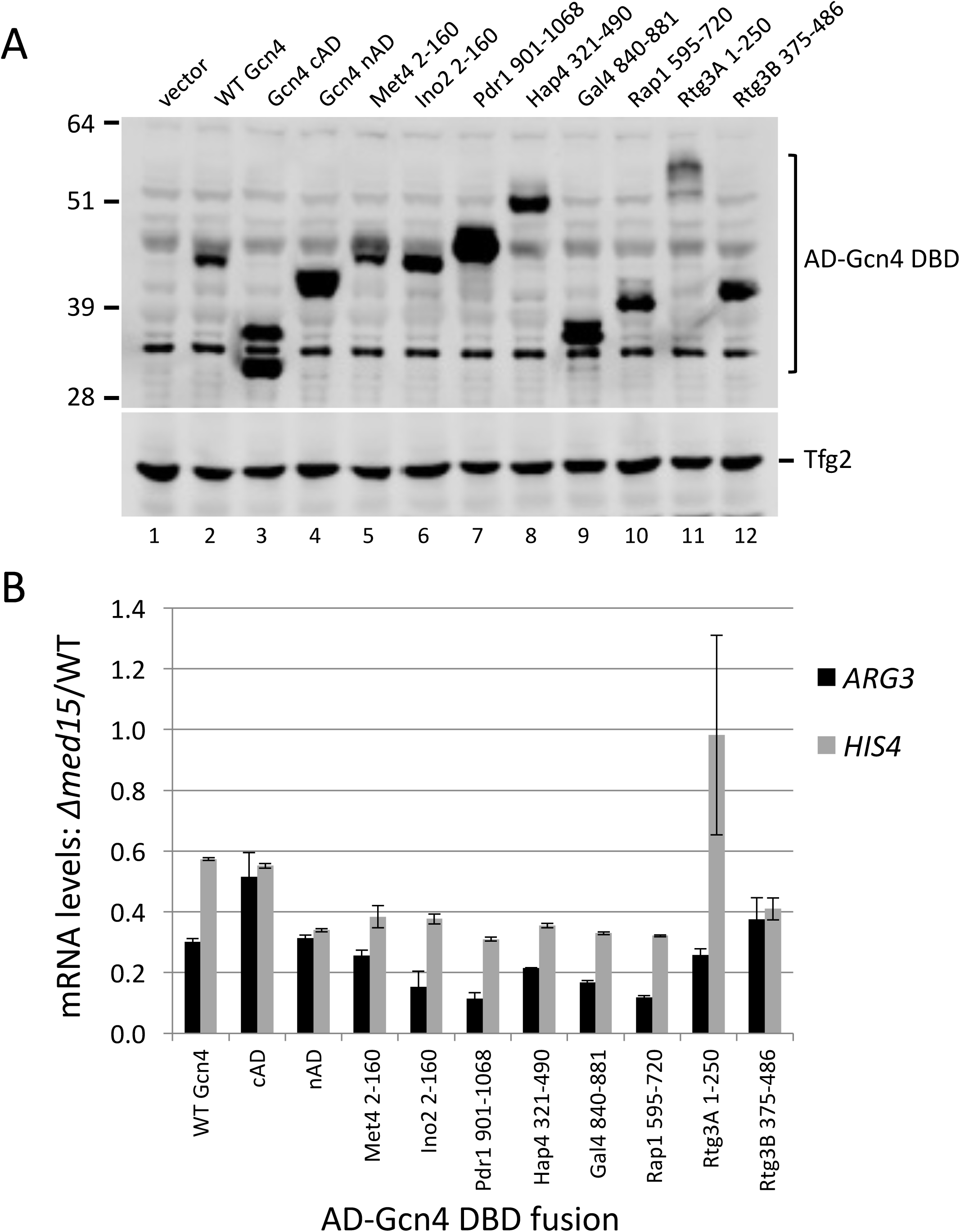
Expression of Gcn4 fusion proteins and Med15-dependence. (**A**) Western blot of whole cell extracts from cells used in Figure 1. Western was probed with anti Gcn4 and anti Tfg2 (TFIIF subunit) as indicated. (**B**). Mediator tail module dependence for the different activators at two Gcn4-responsive genes measured as the ratio of mRNA levels in the *med15Δ/MED15* strains. Data from Figure 1.

TATA-containing Gcn4-activated genes can vary somewhat in their dependence on the Mediator tail module, a direct binding target for Gcn4. For example, *ARG3* shows 5-10-fold dependence on Med15, a Mediator Tail subunit, while *HIS4* shows ~2-fold dependence (27). We measured Mediator tail dependence of these chimeric activators by comparing expression in WT vs *Δmed15.* As previously found with Gcn4, all chimeric ADs showed the strongest Med15 dependence at *ARG3* and somewhat lower dependence at *HIS4* (**Fig 1**; **Fig 2B**) The one outlier among these ADs is the Rtg3 N-terminal AD which showed no Med15 dependence at *HIS4*. From these results, we conclude that nearly all of these ADs function similarly to Gcn4.

### Met4 contains tandem conserved ADs that overlap with ubiquitin-binding domains

Based on in vivo activity, sequence conservation, and previously published work, we focused further characterization on the Met4 and Ino2 ADs. **Fig 3** shows that Met4 residues 65-170 is enriched in both hydrophobic and acidic residues and that it contains tandem 22 residue long sequence blocks that are conserved among closely related yeasts. Both of these conserved regions are predicted to have propensity for alpha helix formation (**Fig 3**).

**Figure 3.**
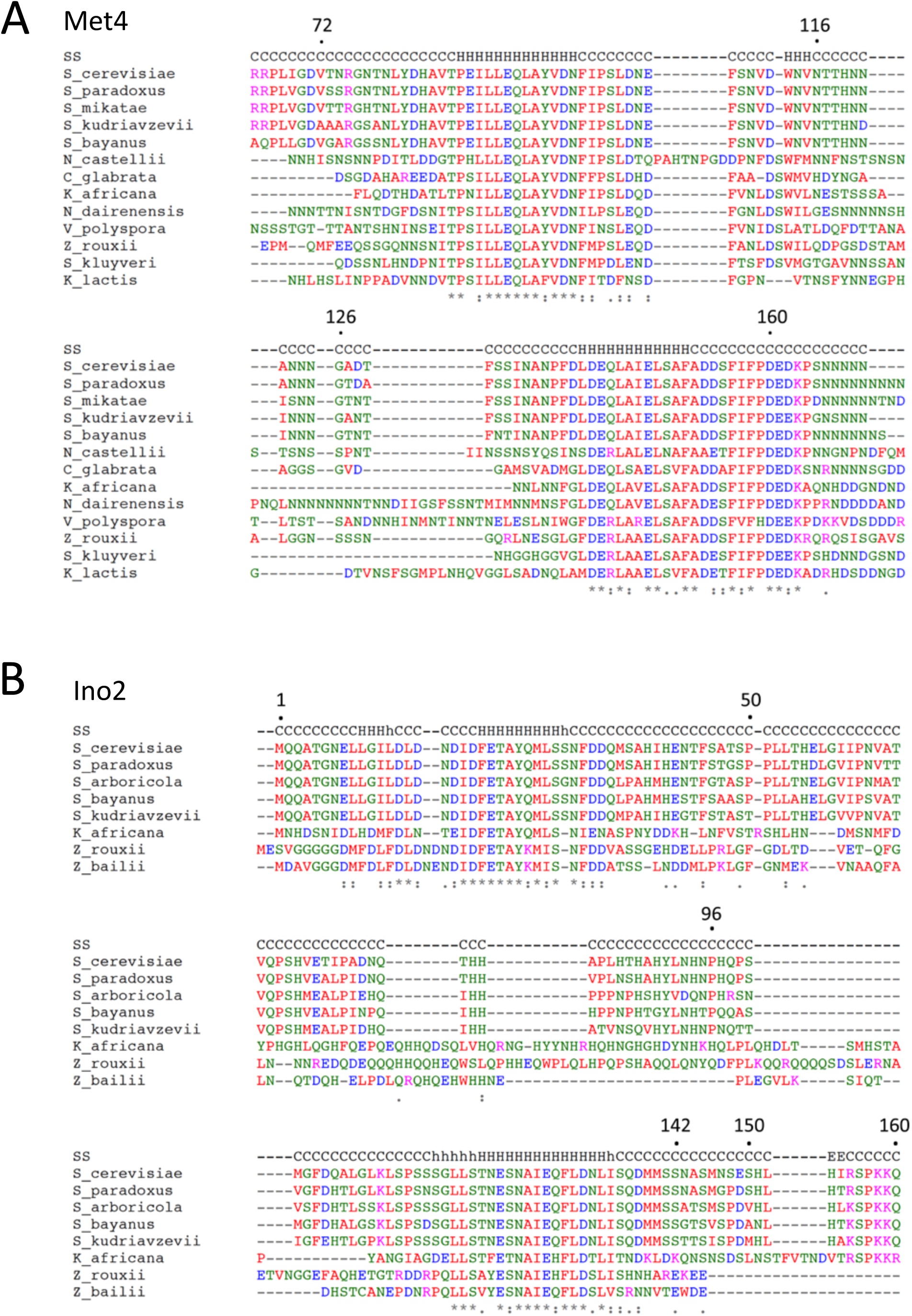
Conservation of activation domains in closely related yeasts. (**A**) Met4 and (**B**) Ino2 sequences aligned by Clustal Omega (57) with secondary structure predictions from Ali2D (58).

Yeast Met4 is a bZIP protein that activates the transcription of at least 45 genes involved in sulfur metabolism (42, 43). Met4 is recruited to regulatory regions by the DNA binding proteins CBF1 and the related factors Met31/32, while cofactor Met28 acts to stabilize these DNA-bound complexes (44, 45). Prior analysis of Met4-LexA fusions showed that Met4 residues 79-160 contains transcription activation function (35). Met4 activator function is known to be regulated by both ubiquitylation and by Ub binding. Met4 is modified by a relatively short poly Ub chain at residue K163 (46), located at the C-terminal edge of the second conserved sequence block. Eliminating ubiquitylation by the mutation K163R activates Met4 similarly to growth in inducing conditions but has little if any effect on protein stability. These findings suggest that Met4-Ub regulates function separately from proteolysis (47, 48). Met4 also contains tandem Ub-binding domains defined by mutations Δ85-96 and Δ135-155 (49). Inactivation of these domains leads to longer Met4 poly Ub chains and decreased protein stability, showing that Ub binding protects Met4 from protein degradation. The Ub-binding domains are contained within the region required for transcription activation and it has not been determined whether these activities are overlapping or independent functions.

A series of deletions was constructed in the Met4-Gcn4 fusion to identify the minimal regions necessary for AD function at *ARG3*, *HIS4* and *ILV6* (**Fig 4**). As with the other chimeric activators above, protein expression levels did not correlate with AD function (**Fig S1**). Consistent with previous observations, Met4 residues 72-160 encode 85-92% of Met4 AD function (35). Further deletions demonstrated that Met4 contains tandem ADs, with the functional regions centered on the two conserved sequence blocks. Met4 72-116 contains 34-62% of Met4 AD function, depending on the activated gene. Met4 126-160 contains 18-55% of Met4 AD function, again depending on the target gene. Because of this gene-specific activation function, the two Met4 ADs synergize at *ARG3* but are approximately additive in activity at *HIS4*. We speculate that this may be due to different coactivator dependencies at these genes.

**Figure 4.**
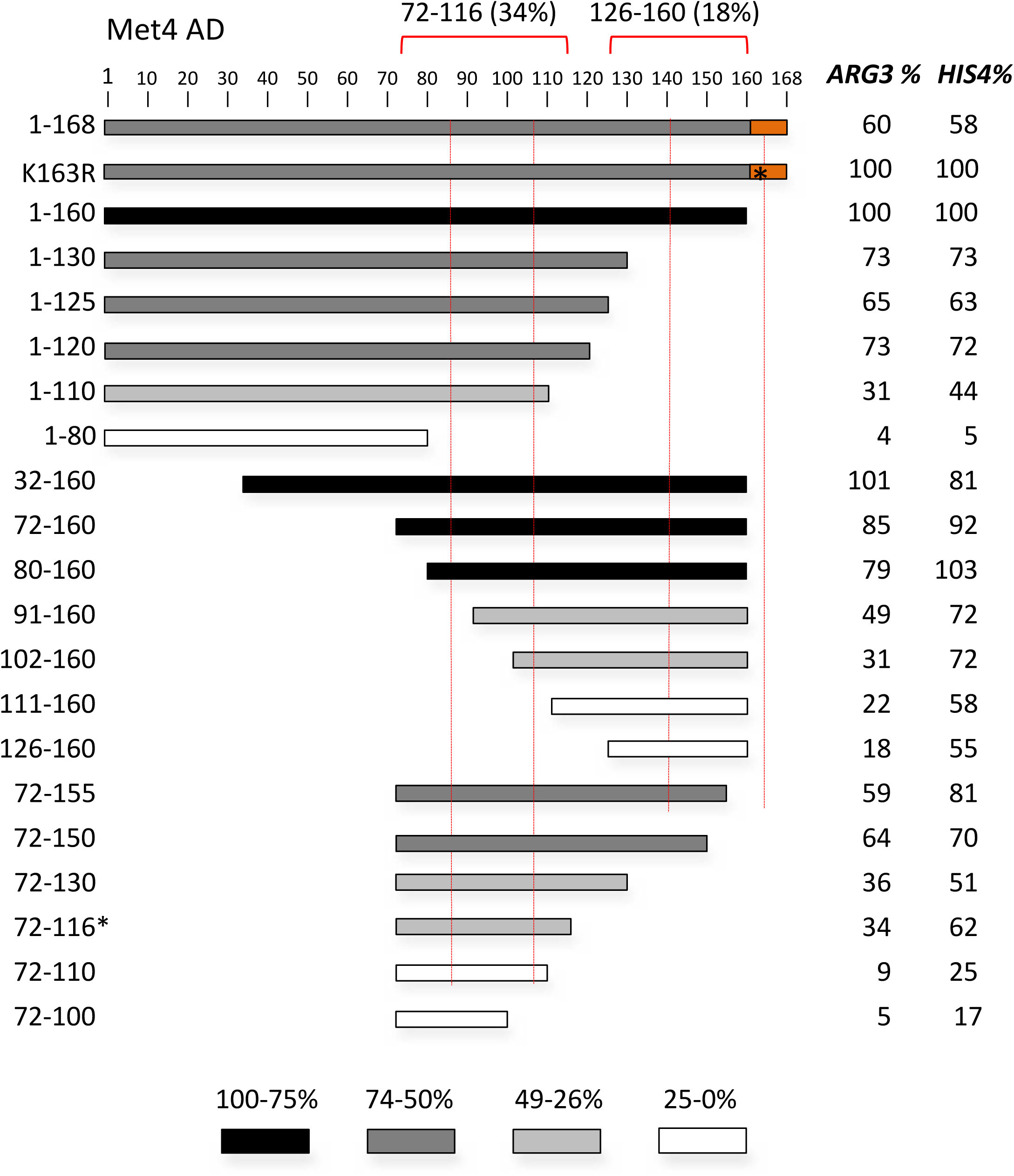
Met4 tandem activation domains. Shown are the Met4 derivatives fused to Gcn4 and assayed for activation of *ARG3*, *HIS4* and *ILV6* as in Figure 1. Protein segments are shaded according to the percent activity compared with 1-160. Red dotted lines indicate the two conserved sequence blocks from Figure 3. The orange block indicates a repressive element and the * indicates the K163R mutation that blocks protein ubiquitylation. Red brackets indicate the limits of the individual ADs at *ARG3* and the percent activity at *ARG3* compared to Met4 1-160.

The deletion analysis also found that Met4 residues 161-168 repress AD function 40-50%. Part of this region is conserved and contains the ubiquitinated residue K163 (47, 48). Western analysis is consistent with modification at this residue as this fusion protein migrates in a series of slower mobility species in SDS PAGE (**Fig S1**), although protein levels appear unchanged compared to Met4 1-160-Gcn4. All derivatives lacking residues 161-168 show no apparent modification. Mutation of K163 to R in the 1-168-Gcn4 construct eliminates both this protein modification and repressive function (**Fig 4; Fig S3**). Unexpectedly, blocking ubiquitination led to lower levels of the fusion protein. This again shows that there is little or no correspondence between protein levels and activation activity in this system.

We next examined the importance of conserved and acidic residues within each Met4 AD for transcription of *ARG3* and *HIS4* (**Fig 5**). For Met4 72-116, alanine substitution at three blocks of conserved hydrophobic residues showed at least a 2-fold decrease in function at one or both of the Gcn4-activated genes. In contrast, mutation of two conserved acidic residues gave no more than a 40% decrease in function. Therefore, like at Gcn4, the hydrophobic residues, not the acidic residues are most important for function. A similar finding was observed with the second AD, Met4 131-160, where three groups of hydrophobic residues are important for function while mutation of two groups of acidic residues showed no major decrease in activity.

**Figure 5.**
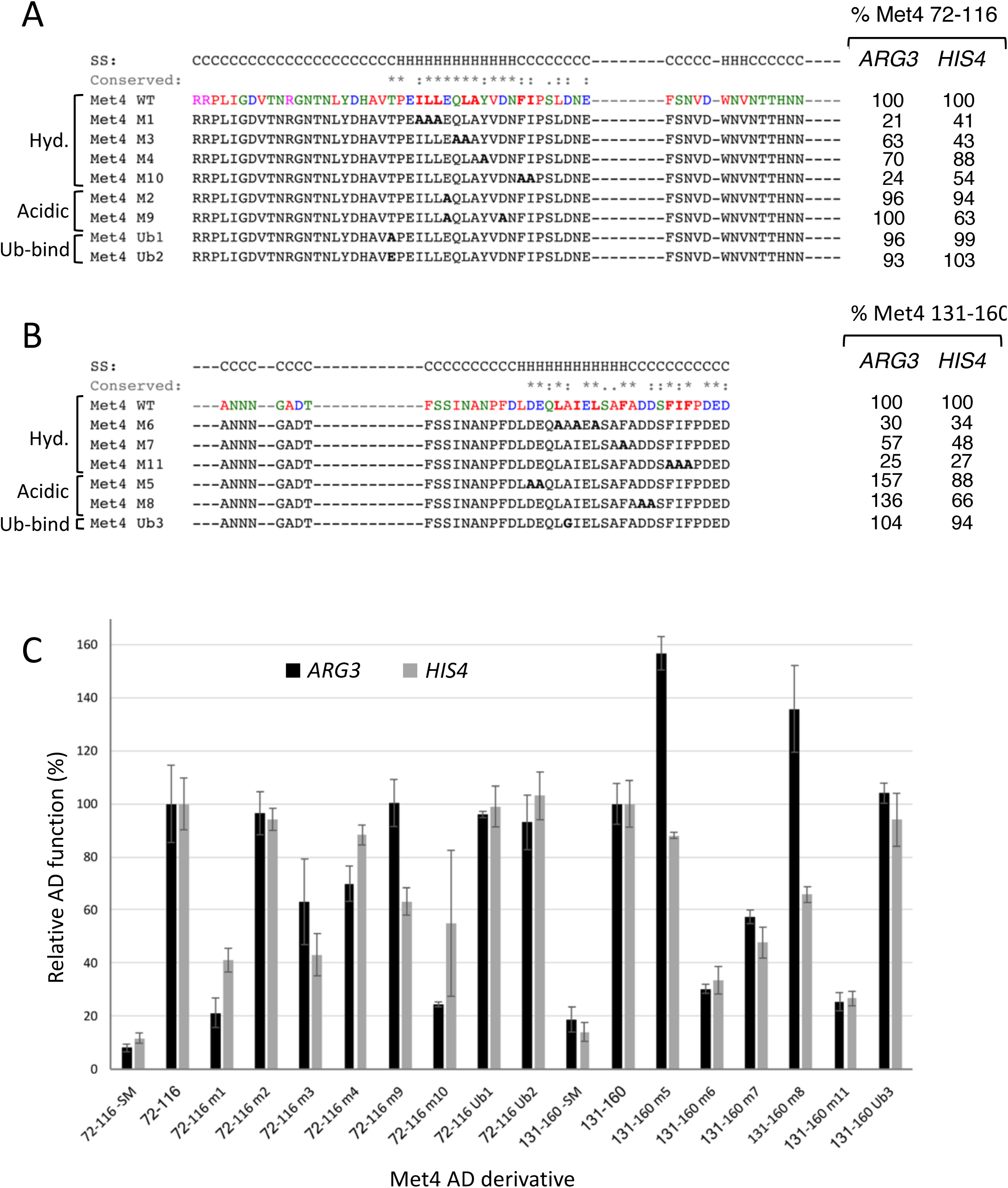
Hydrophobic but not acidic residues are important for Met4 AD function. (**A**) and (**B**) show mutations in the two Met4 ADs that were targeted for Alanine substitution and the resulting effects on induced expression from *ARG3* and *HIS4*. Residues are color coded by amino acid type. Secondary structure predictions and sequence conservation is from Fig 3. (**C**) Quantitation of Met4-Gcn4 fusion protein activity measured by RT qPCR. Data used for (A and B).

As described above, Met4 contains tandem Ub-binding domains that overlap the two conserved sequence blocks in the AD region. To test whether Ub binding and AD function are separable, we tested three mutations that are known to inhibit or eliminate Ub binding (**Fig 5**). Within the 72-116 AD, mutations T86A and T86E, are both known to eliminate Ub binding (An Tyrrell, Karin Flick, and Peter Kaiser, personal communication). These two mutants retained at least 96% wild type function with no major changes in protein level (**Fig S3**). Mutation A145G in the context of residues 131-160, a mutation known to limit Ub binding (47), also caused no decrease in AD function but did significantly reduce fusion protein levels (**Fig S3**). Therefore, we conclude that the Ub binding function of Met4 is not required for activator function, although the two sequences overlap.

### Ino2 contains tandem conserved ADs that require both hydrophobic and acidic residues for function

Next, we examined residues important for Ino2 AD function in the Gcn4 chimeras. Ino2 and Ino4 are bHLH factors required for transcriptional regulation of yeast structural genes involved in phospholipid biosynthesis. Both proteins are required for sequence-specific DNA binding but only Ino2 contains transcription activation function (36, 50). Previous analysis showed that, when fused to the Gal4 DBD, Ino2 residues 1-35 and 98-135 have activator function and were termed AD1 and AD2 (51). Mutagenesis of the AD1 showed that both hydrophobic and acidic residues are required for function. AD2 overlaps with the binding site for the repressor Opi1 (Ino2 residues 118-135), that targets Ino2 to repress transcription in response to high levels of inositol and choline (52). Mutagenesis of AD2 suggested that residues required for Opi1-dependent repression and AD2 function only partially overlap.

Like Met4, the Ino2 AD region contains two blocks of conserved sequence enriched for hydrophobic and acidic residues of 29 and 21 residues that overlap with Ino2 AD1 and AD2 (**Fig 3B**). Both of these conserved regions are predicted to have propensity for alpha helix formation. When fused to the Gcn4 DBD and, in agreement with earlier work, we found that each of the two Ino2 ADs activate *ARG3* and *HIS4* (**Fig 6**). In our system, the minimal segments necessary for AD function are Ino2 residues 1-41 and 96-160. The intervening region between these two regions can be deleted with less than ^~^2-fold decrease in function. Unexpectedly, we found that the C-terminal AD contained a region that repressed function. Deletion of residues 143-150 increased activator function 2-3-fold depending on the promoter assayed (**Fig 6;** orange rectangle). There were no obvious features in the sequence of this region that explain this repressive activity. Although the Ino2 C-terminal AD is reportedly targeted by the Opi1 repressor (52), we found that activity of neither AD was repressed by the addition of inositol and choline (not shown). This is consistent with a report that chimeric Ino2-LexA constructs lacking the Ino2 DBD, are refractory to Opi1 repression (53).

**Figure 6.**
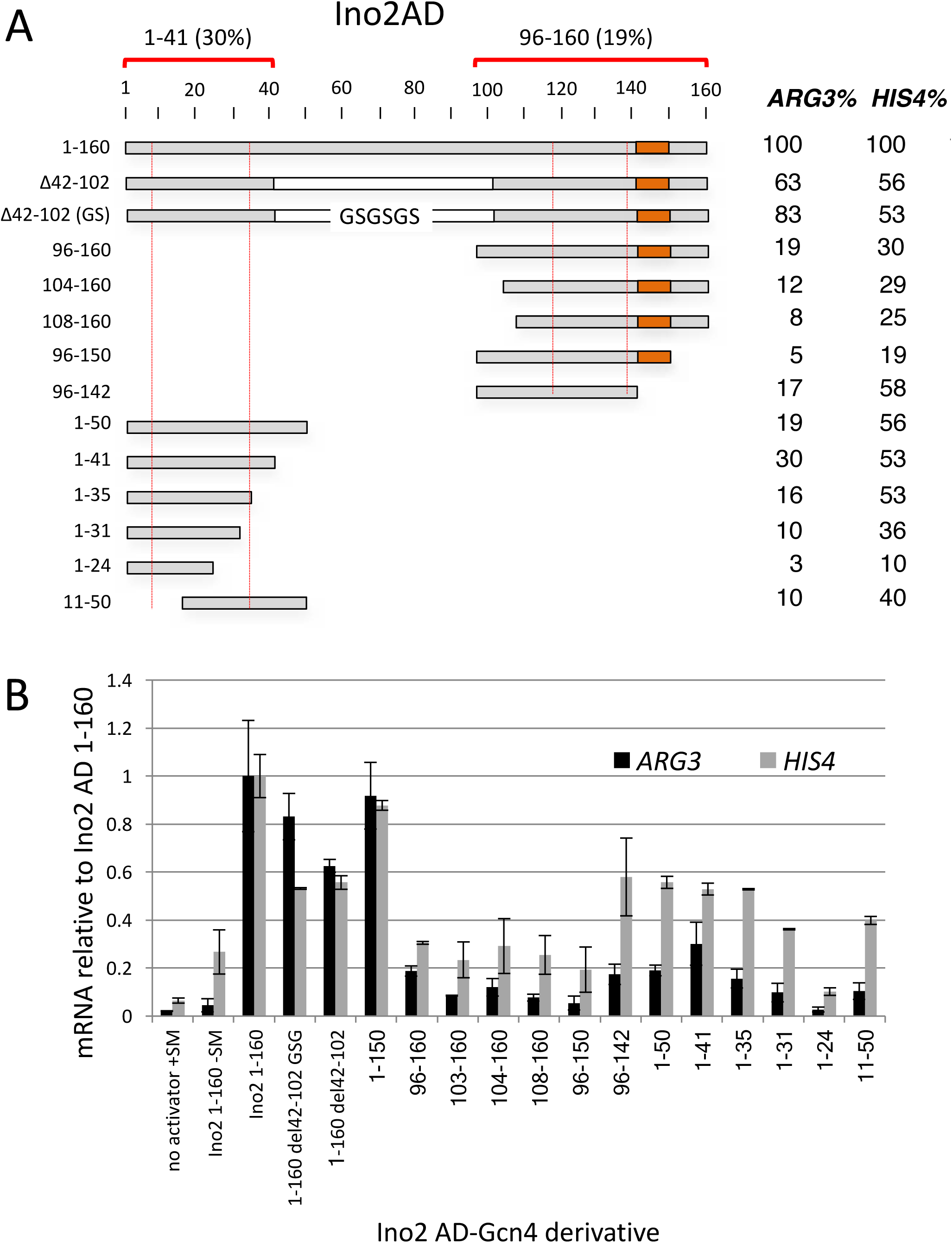
Activity of the Ino2 tandem activation domains. (**A**) Shown are the Ino2 derivatives fused to Gcn4 and assayed for activation of *ARG3* and *HIS4* as in Figure 1. Red dotted lines indicate the two conserved sequence blocks from Figure 3. The orange block indicates an inhibitory element. Red brackets indicate the limits of the two ADs at *ARG3* and the percent activity on *ARG3* compared to Ino2 1-160.

To explore residues important for function of the Ino2 ADs beyond those found in previous work, we mutagenized the individual ADs by double or triple substitution of Ala for hydrophobic, acidic and polar residues. Function was monitored at *ARG3* and *HIS3* under +SM inducing conditions. Residues required for the Ino2 N-terminal AD were distributed over 29 amino acids, almost all of which were in the conserved sequence block (**Fig 7**). We found that 5 sets of hydrophobic mutations reduced activity ~50% or more on at least one Gcn4-dependent gene. In addition, a triple mutation of conserved acidic residues was as detrimental as most mutations of hydrophobic residues. For the C-terminal AD, we found that mutations reducing function were located within a 40-residue segment, much larger than the conserved sequence block (**Fig 7**). Ala substitutions that reduced function by at least 60% on one or both of the Gcn4-dependent genes included 5 sets of hydrophobic residues and one triple mutation of three conserved acidic residues. Unique to this AD, we found that mutation of conserved residues S120, T121 to Ala reduced activity by at least 4-fold. Two mutations of other polar residues did not affect function. In summary, residues important for both Ino2 ADs are distributed over 29-40 residues and include both hydrophobic, and acidic side chains.

**Figure 7.**
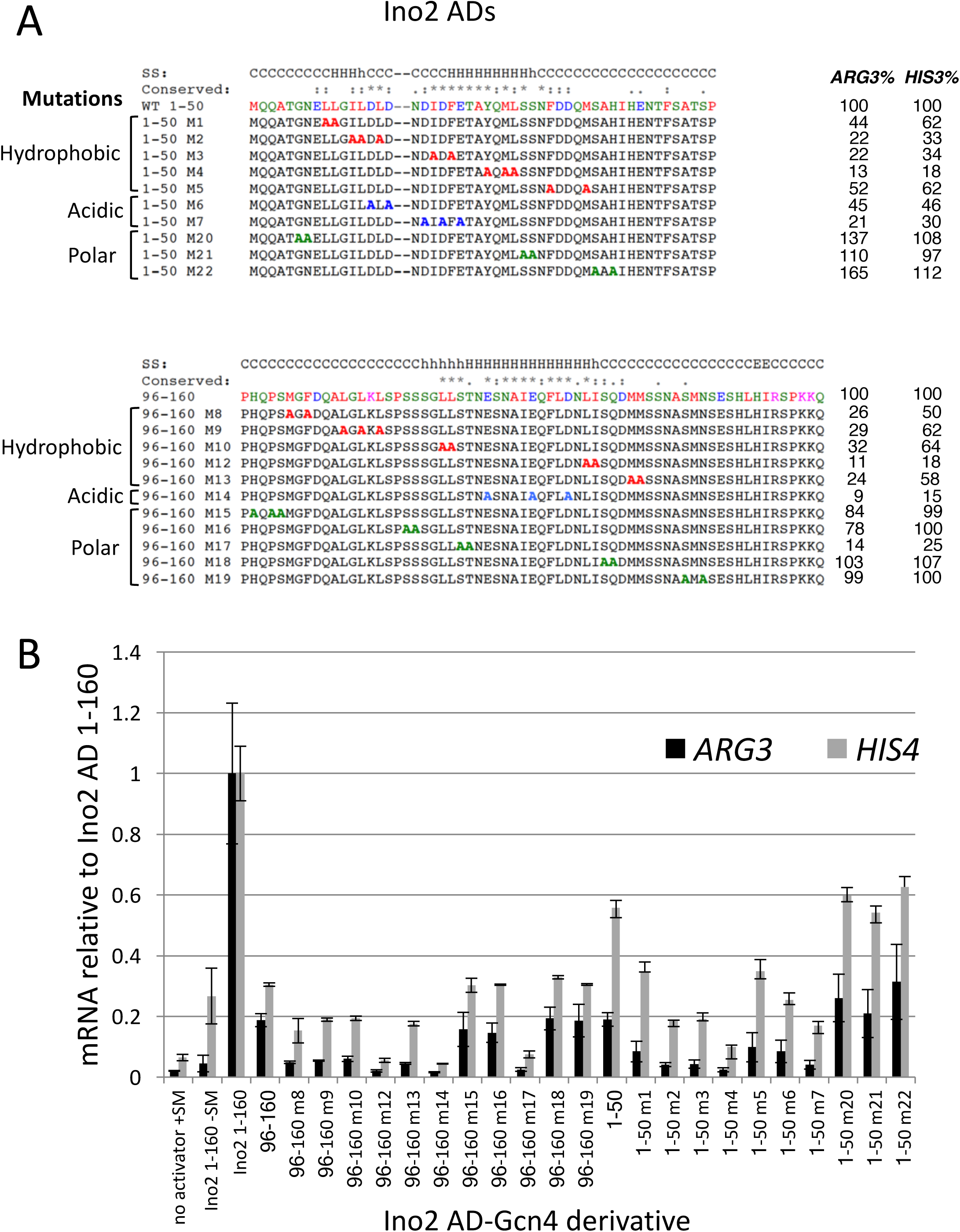
Hydrophobic, acidic, and polar residues are important for Ino2 AD function. (**A**) and (**B**) show mutations in the two Ino2 ADs that were targeted for Alanine substitution and the resulting effects on induced expression from *ARG3* and *HIS4*. Residues are color coded by amino acid type. Secondary structure predictions and sequence conservation is from Fig 3. (**C**) Quantitation of Ino2-Gcn4 fusion protein activity measured by RT qPCR.

### Met4 and Ino2 bind multiple Med15 activator binding domains

To explore the interactions between Med15 and the Ino2 and Met4 tandem ADs, we used purified proteins in combination with isothermal titration calorimetry (ITC) and/or fluorescence polarization (FP) to measure the affinities and thermodynamic properties of these interactions (**Figures 8-10**; summarized in **Table 1**). Binding between the ADs and the individual Med15 activator binding domains was monitored using ITC. We were not able to use ITC to monitor binding to the longer Med15 polypeptides (KIX +ABD1,2,3 and ABD1,2,3) so FP was employed to monitor N-terminal fluorescently-labeled AD peptides binding to Med15. For comparison of the two methods we used FP and ITC to monitor AD binding to ABD3 and the results were similar. Met4 affinities monitored by either approach were within 20% and Ino2 affinities were within ^~^3-fold. For the discussion below, the affinities are compared using the ITC values where available.

**Figure 8.**
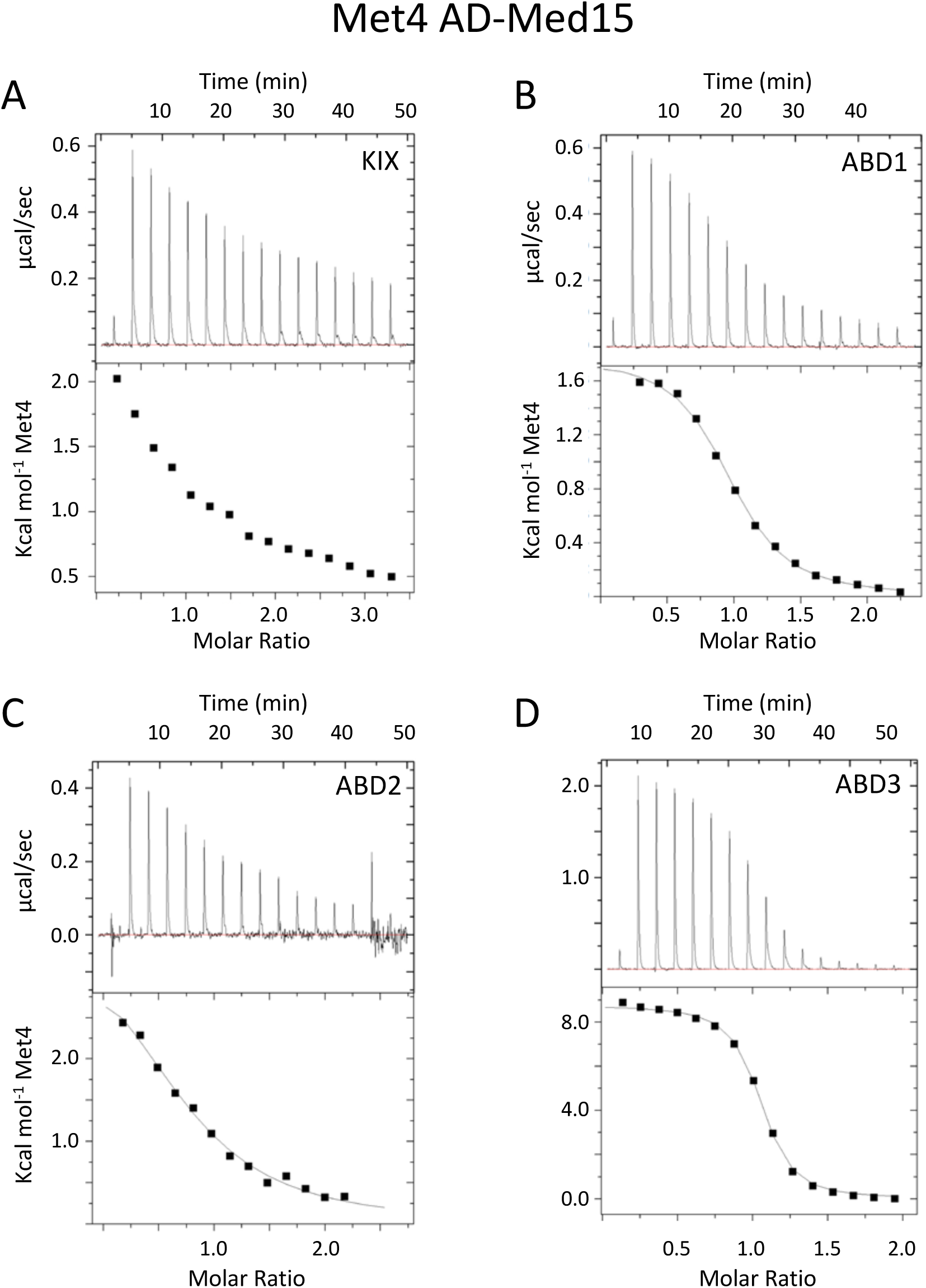
Measurement of Met4-Med15 binding by Isothermal titration calorimetry. ITC was used to determine the affinity and thermodynamic parameters of Met4 72-160 interactions with the Med15 KIX domain (**A**), Med15 ABD1 (**B**), Med15 ABD2 (**C**), and Med15 ABD3 (**D**). All assays were performed and curves were fit as described in Materials and Methods.

**Figure 9.**
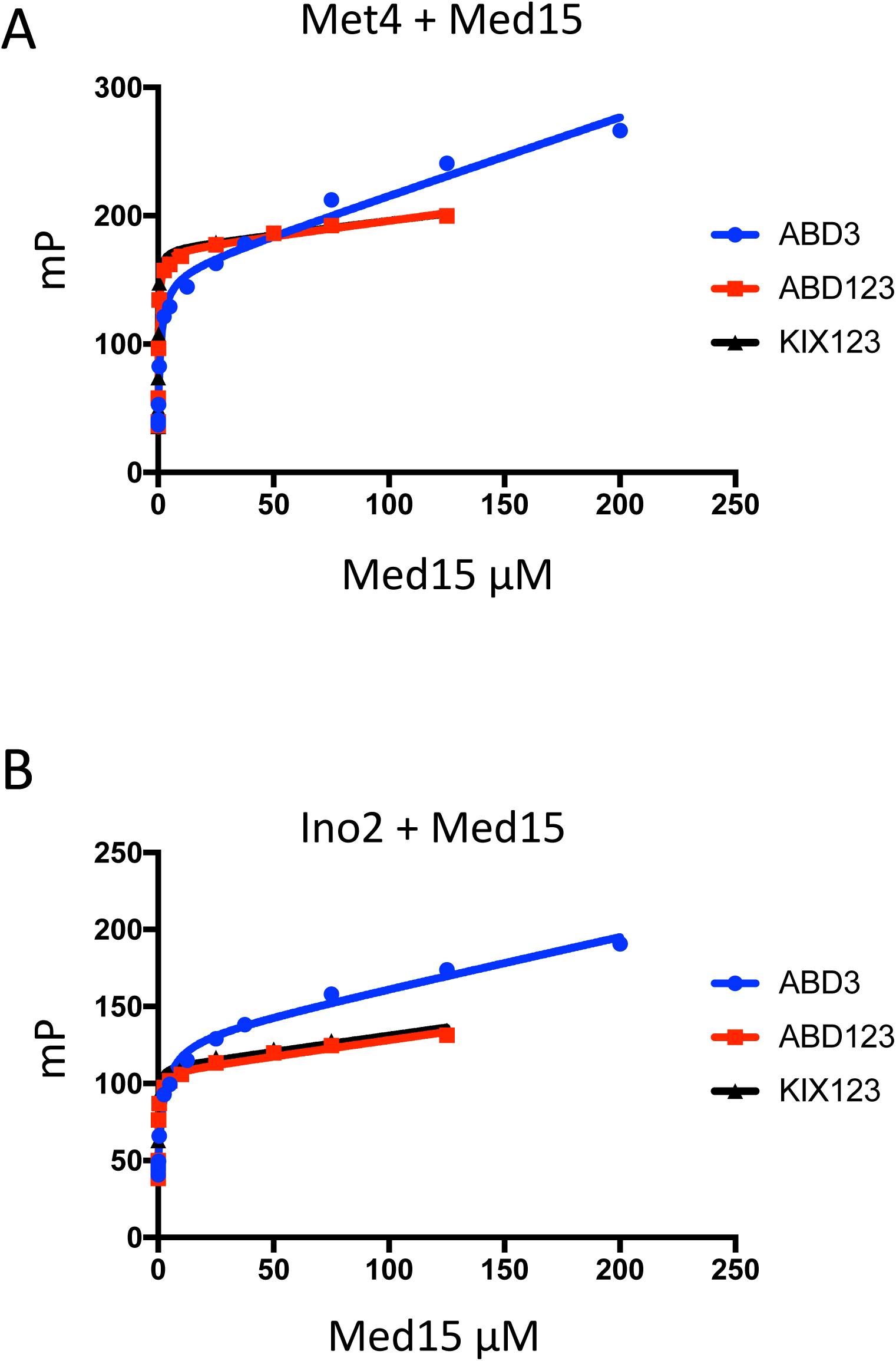
Measurement of activator-Med15 binding by Fluorescence Polarization. FP was used to assay binding of Oregon Green-labeled Met4 72-160 (**A**) or Ino2 1-41-(GS)_3_-96-160 (**B**) to Med15 ABD3, Med15 ABD123, and Med15 KIX+ ABD123. All assays were performed in triplicate, curves were fit as described in Materials and Methods.

**Figure 10.**
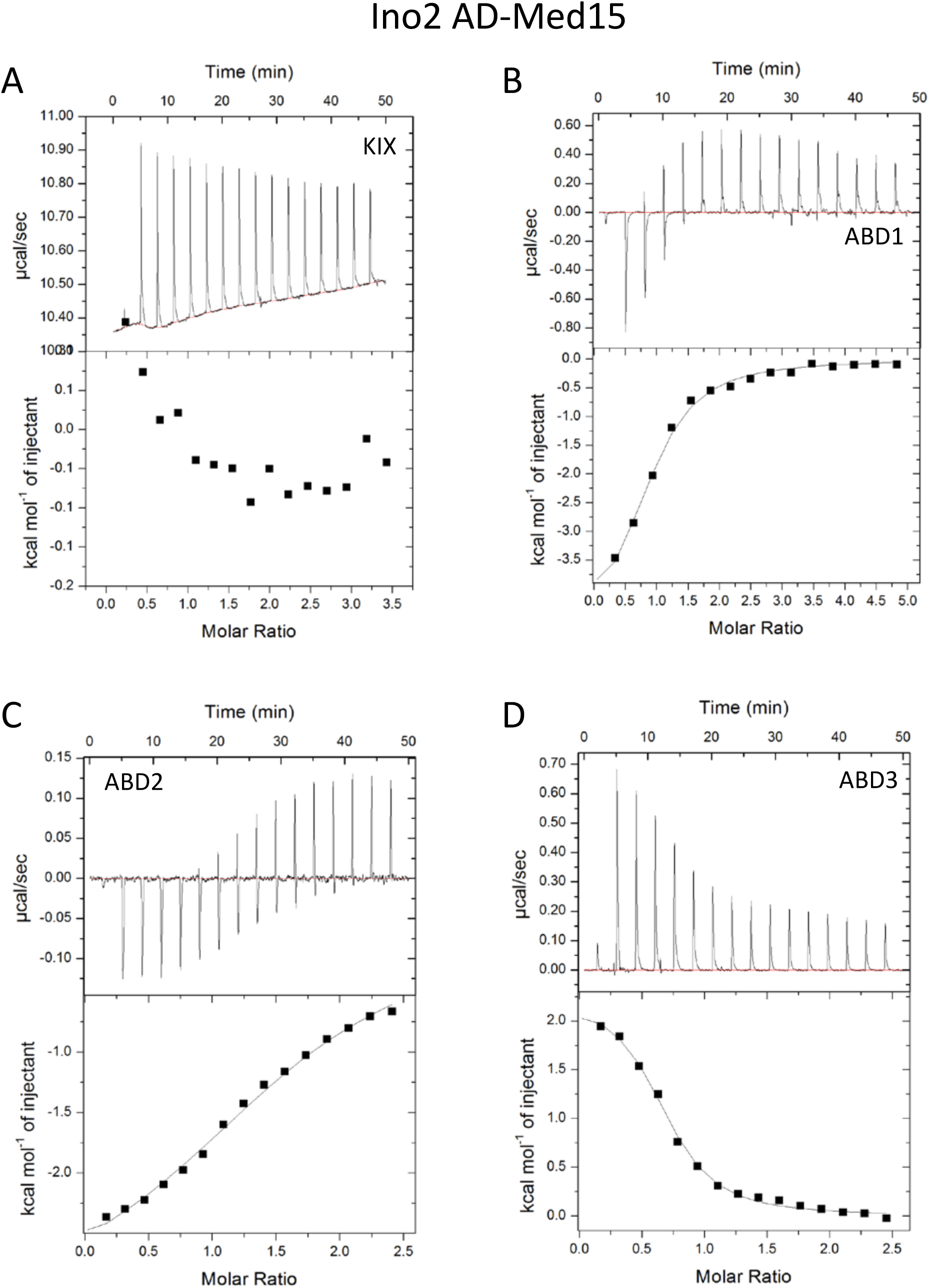
Measurement of Ino2-Med15 binding. ITC was used to determine the affinity and thermodynamic parameters of Ino2 1-41-(GS)_3_-96-160 interactions with the Med15 KIX domain (**A**), Med15 ABD1 (**B**), Med15 ABD2 (**C**), and Med15 ABD3 (**D**). All assays were performed and curves were fit as described in Materials and Methods.

**Table 1.** Affinity of Met4 and Ino2 ADs binding to Med15 derivatives. (**A**) Affinity of Met4-Med15 interactions. ITC: isothermal titration calorimetry; FP: fluorescence polarization. For ITC measurements, calculated values of ΔH cal/mole (enthalpy), ΔS cal/mole/deg (entropy) and N (molar ratio) are given. NM = not measurable; N/A not applicable. Proteins used are Met4: 72-160; Med15 KIX: 6-90; Med15 ABD1: 158-238; Med15 ABD2: 277-368; Med15 ABD3: 484-651; Med15 ABD1,2,3; 158-651 Δ239-272, Δ373-483; Med15 KIX+ABD1,2,3: 1-651 Δ239-272, Δ373-483. (**B**) Affinity of Ino2-Med15 interactions. Same nomenclature as in (A). Proteins used are Ino2: 1- 41-(GS)_3_-96-160; Med15 KIX: 6-90; Med15 ABD1: 158-238; Med15 ABD2: 277-368; Med15 ABD3: 484-651; Med15 ABD1,2,3; 158-651 Δ239-272, Δ373-483; Med15 KIX+ABD1,2,3: 1-651 Δ239-272, Δ373-483.

| Med15 | Kd (μM) | ΔH | ΔS | N | Method |
| --- | --- | --- | --- | --- | --- |
| KIX | NM | N/A | N/A | N/A | ITC |
| ABD1 | $5.9 \pm 0.5$ | $1780 \pm 26$ | 30 | 0.85 | ITC |
| ABD2 | $20.4 \pm 3.4$ | $3789 \pm 335$ | 34.3 | 0.80 | ITC |
| ABD3 | $1.36 \pm 0.1$ | $8920 \pm 46$ | 57 | 1.01 | ITC |
| ABD3 | $1.11 \pm 0.23$ | N/A | N/A | N/A | FP |
| ABD1,2,3 | $0.283 \pm 0.027$ | N/A | N/A | N/A | FP |
| KIX+ABD1,2,3 | $0.196 \pm 0.015$ | N/A | N/A | N/A | FP |

**(B) Affinity of Ino2-Med15 interactions**
| Med15 | Kd (μM) | ΔH | ΔS | N | Method |
| --- | --- | --- | --- | --- | --- |
| KIX | NM | N/A | N/A | N/A | ITC |
| ABD1 | $25.3 \pm 3.4$ | $-4916 \pm 286$ | 4.4 | 0.85 | ITC |
| ABD2 | $33.8 \pm 3.6$ | $-2440 \pm 73$ | 12.2 | 1.48 | ITC |
| ABD3 | $7.75 \pm 0.9$ | $2250 \pm 61$ | 31 | 0.66 | ITC |
| ABD3 | $2.33 \pm 0.4$ | N/A | N/A | N/A | FP |
| ABD1,2,3 | $0.314 \pm 0.051$ | N/A | N/A | N/A | FP |
| KIX+ABD1,2,3 | $0.213 \pm 0.019$ | N/A | N/A | N/A | FP |

For both ADs, we were unable to detect binding to the Med15 KIX domain. This behavior is identical to that of the Gcn4 ADs (27). In contrast, both ADs bound to ABD1, 2, and 3. For Met4, the order of highest to lowest binding was ABD3>ABD1>ABD2 with affinities ranging from 1 to 20 micromolar. The relative order of Ino2 interactions was the same, but all of the individual interactions were weaker compared to Met4, ranging from 8-34 micromolar. Highest affinity interactions were with Med15 polypeptides containing all ABDs: KIX + ABD1,2,3 and ABD1,2,3. For both activators, constructs containing the KIX domain had the highest affinity for the ADs even though binding to the isolated KIX domain was undetectable in our assays. This is consistent with results found for Gcn4, where the KIX domains seemed as functionally important as any of the Med15 ABDs (27) and where KIX + ABD1,2,3 had the highest affinity for the tandem Gcn4 ADs (20). Combined, our results show that Med15 polypeptides with multiple ABDs have much higher affinity for Met4 and Ino2. For example, Met4 binds KIX + ABD1,2,3 with ^~^7-fold higher affinity compared with ABD3 (Kd of 0.196 versus 1.36 micromolar) and Ino2 binds KIX + ABD1,2,3 with 36-fold higher affinity compared with ABD3 (Kd of 0.21 vs 7.8 micromolar). Finally, despite our finding that the Met4 had higher affinity than Ino2 for the individual Med15 ABDs, the affinity of Ino2 and Met4 for the longer Med15 polypeptides was remarkably similar (Kd ^~^0.2 micromolar for KIX + ABD1,2,3 and ^~^0.3 micromolar for ABD1,2,3).

Our previous work showed different thermodynamic behavior in the mechanism of Med15 binding to the two Gcn4 ADs (6). For example, the Gcn4 central activation domain (cAD) binding to ABD1 is exothermic with a favorable change in enthalpy and a small but positive entropy change. In contrast, the Gcn4 nAD binding to the individual Med15 ABD1, 2, and 3 domains are endothermic, with unfavorable changes in enthalpy counteracted by large positive changes in entropy. Binding of Met4 and Ino2 ADs also showed surprising and varied thermodynamic behavior depending on the combination of activator and Med15 ABD (**Figs 8, 10 and Table 1**). For example, binding of Gcn4 nAD, Met4 AD and Ino2 AD to ABD3 is consistently endothermic. In contrast, binding to ABD1 and ABD2 can be endo or exothermic depending on the activator.

### Crosslinking reveals heterogeneous AD-ABD interactions within the Met4-Med15 complex

The individual binding measurements above showed that Met4 and Ino2 interact with both the individual ABDs and longer Med15 polypeptides but these experiments cannot show whether the relative affinity or ABD specificity changes in the larger complexes. For example, these studies show that the KIX domain contributes to overall affinity, but does not answer whether there is a direct contact between the AD and KIX. To examine the binding mechanism of the Met4 tandem ADs with the full-length Med15 activator-binding regions, we used the crosslinker EDC, which crosslinks acidic side chains to lysine (**Fig 11**; **Table S2**). EDC is a zero-length crosslinker, linking only closely positioned residues and leaving no linker in the crosslinked product. Analysis of the crosslinked products by mass spectrometry identified crosslinks between the individual Met4 ADs and Med15 KIX, ABD1, ABD2 and ABD3. All of the intermolecular crosslinks were between acidic residues in Met4 and lysine residues in Med15. Surprisingly, fewer crosslinks were detected with ABD3, which individually has the highest affinity for Met4 compared to the other ABDs. Our combined results show that Met4 makes direct contacts with KIX and that there is no unique protein complex formed upon binding of Met4 to Med15. Rather, our results are consistent with the tandem ADs rapidly sampling the Med15 ABDs in a large dynamic fuzzy complex as previously proposed (6).

**Figure 11.**
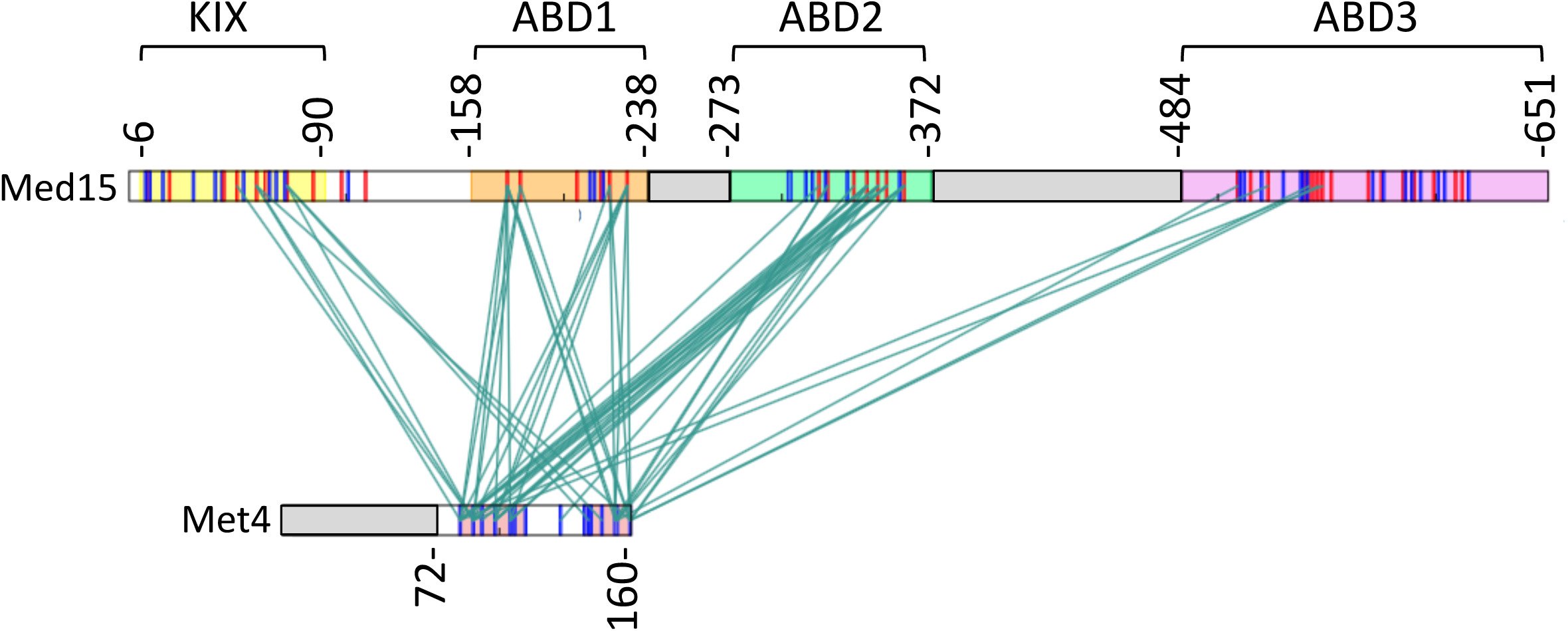
Met4 ADs interacts via a heterogeneous complex with the three ABDs of Med15. Mass spectrometry crosslinking experiments show crosslinks are formed between regions throughout Met4 AD and each of the Med15 ABD regions and to KIX. Crosslinks between Met4 72-160 and Med15 KIX123 are shown in the context of Met4 1-160 and Med15 1-651. Deleted regions are indicated by the grey boxes. Red bars indicate lysine residues. Blue bars indicate aspartic acid and glutamic acid residues. Conserved regions of the Met4 AD are shaded pink. Regions of Med15 containing the ABDs are colored as follows: KIX (aa 6-90), yellow; ABD1 (aa 158-238), orange; ABD2 (aa 272-372), green; ABD3 (aa 484-651), purple.

## Discussion

Compared with most protein-protein interactions, interactions of transcription activators with their targets are unusual. The primary sequence of ADs is not obviously conserved among different activators, the factors are intrinsically disordered and they interact with multiple distinct targets having no obvious similarity. However, these properties undoubtedly allow many activators to function through a variety of coactivators and to modulate transcription at varied promoters with different coactivator requirements. Here, we have focused our investigations on characterizing two strong yeast activators, Met4 and Ino2, to identify common and distinct features of yeast ADs. Examining Met 4, Ino2, and 7 other strong yeast activation domains, we found that all but one has similar dependence on the Mediator Tail module subunit Med15 for activation of two TATA-containing reporter genes.

Like Gcn4, both Met4 and Ino2 have tandem ADs that are enriched for acidic and hydrophobic residues. Tandem ADs may be another feature common to strong activators in eukaryotes, as mammalian viral and human activators such as VP16 and p53 also have tandem ADs. For Met4 and Ino2, these individual ADs were identified both functionally and as blocks of moderately conserved sequences with helical propensity imbedded in non-conserved flanking sequences. This, along with previous work shows that, although ADs do not have a common primary sequence motif, there are specific sequence requirements that constitute a functional AD.

Both individual ADs of Met4 are of intermediate length: the conserved sequence blocks are 22 residues long for both and mutagenesis of conserved residues shows that conserved sequences of 13 and 15 residues long are required for most of the AD function. Mutagenesis of the ADs found that only hydrophobic residues were critical for normal function – identical to the finding of critical hydrophobic but not acidic residues in the Gcn4 ADs (18, 19, 24). The individual Ino2 ADs are larger than Met4 ADs with 29 and 21 residue conserved sequence blocks. Mutagenesis of the N-terminal AD found that a stretch of 29 residues was required for maximum function that almost precisely coincided with the conserved sequence block. However, the C-terminal AD was larger with functionally important amino acids distributed over a span of 40 residues.

We also found that both Ino2 ADs contained functionally important hydrophobic and acidic residues. The acidic residues may function through non-specific electrostatic interactions with the coactivator targets or alternatively may make direct and specific contacts.

Monitoring the binding of Ino2 and Met4 to Med15 showed that they behaved in many respects like Gcn4. All bind Med15 ABD1, ABD2, and ABD3 with micromolar affinity and binding to the Med15 KIX domain is undetectable in our assays. Binding of the tandem ADs to larger Med15 polypeptides all have much higher affinity compared to the individual ABDs and the KIX domain contributes to overall affinity under these conditions. These biochemical findings are consistent with our earlier study that showed the normal in vivo response to Gcn4 activation requires multiple Med15 ABDs and the KIX domain. It seems likely that, since these individual binding interactions are weak, multiple binding sites are required to increase the affinity and specificity into a biologically meaningful range (54, 55).

An unexpected observation with Gcn4, Met4 and Ino2 binding to Med15 is that interactions with the individual ABDs could be either exo or endothermic. The endothermic interactions all have large unfavorable changes in enthalpy and are driven by large positive changes in entropy. This behavior is opposite from that expected because of the entropic penalty paid upon binding of a disordered protein. However, it has been proposed that, even in the bound state, IDPs can retain conformational entropy due to “fuzzy” protein interfaces and conformational flexibility of the protein region not in direct contact with the binding partner (56). However, these mechanisms do not seem to fully explain the large, positive entropy changes observed. At this time, we do not understand the mechanism for the large increase in entropy upon binding, but it seems to be ABD and activator-specific and is likely to at least partially result from release of solvent during binding. As an example of thermodynamic specificity, binding of ABD3 to Gcn4, Met4 and Ino2 is endothermic while the thermodynamics of binding to ABD1 and ABD2 is activator-specific. Understanding the mechanism of endothermic binding will be important for not only understanding activator mechanisms and specificity but more generally as a mechanism likely used for molecular recognition by other disordered proteins.

Finally, the Met4-Med15 crosslinking experiments allowed us to probe larger and more physiologically relevant complexes. Upon mixing the Met4 tandem ADs with KIX + ABD1,2,3, crosslinking revealed that the individual Met4 ADs directly interact with each of the Med15 structured domains. This shows that there is no unique Met4-Med15 protein complex and is consistent with the model that multiple Gcn4 ADs rapidly sample individual Med15 ABDs in a large dynamic fuzzy complex (20). Since this crosslinking behavior is identical to that observed with Gcn4, and because Met4 and Ino2 have generally similar properties, we think it likely that all three activators function by similar mechanisms. In the future, it will be important to understand more about both the biochemical properties of these interactions, and how often eukaryotic activators use this mode of protein-protein interaction.

## Author contributions

DP, LW, MB, HR, and SH performed all experimental work. DP purified proteins, measured binding affinities and, along with MB, performed and analyzed crosslinking reactions. LW, HR, and SH created fusion proteins and derivatives and LW and HR analyzed activator function and protein expression. JL and JR performed crosslinking-MS analysis. DP, LW, and SH wrote the paper.

## Acknowledgements

We thank Johannes Soeding and Eckhardt Guthöhrlein (Max Plank, Göttingen and the Gene Center, Munich) for many insightful discussions and analysis of the Ino2 and Met4 AD sequences, Peter Kaiser (UC Irvine) for discussions and permission to cite unpublished work, Laurent Kuras (CEA, CNRS, Université Paris Sud) for discussions, Members of the Hahn lab for assistance and comments during the course of this work, and Lisa Tuttle for comments on the manuscript. Funded by grant from UW School of Medicine to HR and MB, 2P50GM076547 and 5R01GM110064 to JR, and NIH grant 5RO1GM075114 to SH. HR and MB were affiliated with the University of Washington Medical Student Research Training Program.

---

**Table S1.** Strains and Plasmids used in this work.

**Table S2.** Summary of EDC crosslinks within the Met4 72-160 - Med15 KIX123 complex

**Figure S1.** Protein expression of Met4-Gcn4 derivatives. Shown are Western blots analyzing whole cell extracts of cells used for the RT qPCR assays. Blots were probed with anti FLAG or Tfg2 (TFIIF subunit) as indicated.

**Figure S2.** Protein expression of Ino2-Gcn4 derivatives. Shown are Western blots analyzing whole cell extracts of cells used for the RT qPCR assays. Blots were probed with anti FLAG.

**Figure S3.** Protein expression of Ino2 and Met4-Gcn4 derivatives. Shown are Western blots analyzing whole cell extracts of cells used for the RT qPCR assays. Blots were probed with anti Gcn4 and anti Tfg2.

## References

1. Spitz F, Furlong EEM. 2012. Transcription factors: from enhancer binding to developmental control. Nat Rev Genet 13:613–626.

2. Levine M, Cattoglio C, Tjian R. 2014. Looping back to leap forward: transcription enters a new era. Cell 157:13–25.

3. Hahn S, Young ET. 2011. Transcriptional regulation in Saccharomyces cerevisiae: transcription factor regulation and function, mechanisms of initiation, and roles of activators and coactivators. Genetics 189:705–736.

4. Weake VM, Workman JL. 2010. Inducible gene expression: diverse regulatory mechanisms. Nat Rev Genet 11:426–437.

5. Ptashne M, Gann AA. 1990. Activators and targets. Nature 346:329–331.

6. Brzovic PS, Heikaus CC, Kisselev L, Vernon R, Herbig E, Pacheco D, Warfield L, Littlefield P, Baker D, Klevit RE, Hahn S. 2011. The acidic transcription activator Gcn4 binds the mediator subunit Gal11/Med15 using a simple protein interface forming a fuzzy complex. Mol Cell 44:942–953.

7. Feng H, Jenkins LMM, Durell SR, Hayashi R, Mazur SJ, Cherry S, Tropea JE, Miller M, Wlodawer A, Appella E, Bai Y. 2009. Structural basis for p300 Taz2-p53 TAD1 binding and modulation by phosphorylation. Structure/Folding and Design 17:202–210.

8. Kussie PH, Gorina S, Marechal V, Elenbaas B, Moreau J, Levine AJ, Pavletich NP. 1996. Structure of the MDM2 oncoprotein bound to the p53 tumor suppressor transactivation domain. Science 274:948–953.

9. Sigler PB. 1988. Transcriptional activation. Acid blobs and negative noodles. Nature.

10. Uesugi M, Nyanguile O, Lu H, Levine AJ, Verdine GL. 1997. Induced alpha helix in the VP16 activation domain upon binding to a human TAF. Science 277:1310–1313.

11. Stein A, Pache RA, Bernadó P, Pons M, Aloy P. 2009. Dynamic interactions of proteins in complex networks: a more structured view. FEBS J 276:5390–5405.

12. Nguyen Ba AN, Yeh BJ, van Dyk D, Davidson AR, Andrews BJ, Weiss EL, Moses AM. 2012. Proteome-wide discovery of evolutionary conserved sequences in disordered regions. Science Signaling 5:rs1.

13. Das RK, Mao AH, Pappu RV. 2012. Unmasking functional motifs within disordered regions of proteins. Science Signaling 5:pe17.

14. Gould CM, Diella F, Via A, Puntervoll P, Gemünd C, Chabanis-Davidson S, Michael S, Sayadi A, Bryne JC, Chica C, Seiler M, Davey NE, Haslam N, Weatheritt RJ, Budd A, Hughes T, Pas J, Rychlewski L, Trave G, Aasland R, Helmer-Citterich M, Linding R, Gibson TJ. 2010. ELM: the status of the 2010 eukaryotic linear motif resource. Nucleic Acids Res 38:D167–80.

15. Mitchell PJ, Tjian R. 1989. Transcriptional regulation in mammalian cells by sequence-specific DNA binding proteins. Science 245:371–378.

16. Ptashne M, Gann A. 1997. Transcriptional activation by recruitment. Nature 386:569–577.

17. Tompa P. 2002. Intrinsically unstructured proteins. Trends in Biochemical Sciences 27:527–533.

18. Warfield L, Tuttle LM, Pacheco D, Klevit RE, Hahn S. 2014. A sequence-specific transcription activator motif and powerful synthetic variants that bind Mediator using a fuzzy protein interface. Proceedings of the National Academy of Sciences 111:E3506–13.

19. Drysdale CM, Dueñas E, Jackson BM, Reusser U, Braus GH, Hinnebusch AG. 1995. The transcriptional activator GCN4 contains multiple activation domains that are critically dependent on hydrophobic amino acids. Mol Cell Biol 15:1220–1233.

20. Tuttle LM, Pacheco D, Warfield L, Luo J, Ranish J, Hahn S, Klevit RE. 2017. Transcription activator-coactivator specificity is mediated by a large and dynamic fuzzy protein-protein complex. bioRxiv 1–44. DOI: 10.1101-221747.

21. Erkina TY, Erkine AM. 2016. Nucleosome distortion as a possible mechanism of transcription activation domain function. Epigenetics & chromatin 9:40.

22. Qiu H, Chereji RV, Hu C, Cole HA, Rawal Y, Clark DJ, Hinnebusch AG. 2016. Genome-wide cooperation by HAT Gcn5, remodeler SWI/SNF, and chaperone Ydj1 in promoter nucleosome eviction and transcriptional activation. Genome Res 26:211–225.

23. Mittal N, Guimaraes JC, Gross T, Schmidt A, Vina-Vilaseca A, Nedialkova DD, Aeschimann F, Leidel SA, Spang A, Zavolan M. 2017. The Gcn4 transcription factor reduces protein synthesis capacity and extends yeast lifespan. Nat Commun 8:457.

24. Jackson BM, Drysdale CM, Natarajan K, Hinnebusch AG. 1996. Identification of seven hydrophobic clusters in GCN4 making redundant contributions to transcriptional activation. Mol Cell Biol 16:5557–5571.

25. Brown CE, Howe L, Sousa K, Alley SC, Carrozza MJ, Tan S, Workman JL. 2001. Recruitment of HAT complexes by direct activator interactions with the ATM-related Tra1 subunit. Science 292:2333–2337.

26. Fishburn J, Mohibullah N, Hahn S. 2005. Function of a eukaryotic transcription activator during the transcription cycle. Mol Cell 18:369–378.

27. Herbig E, Warfield L, Fish L, Fishburn J, Knutson BA, Moorefield B, Pacheco D, Hahn S. 2010. Mechanism of Mediator recruitment by tandem Gcn4 activation domains and three Gal11 activator-binding domains. Mol Cell Biol 30:2376–2390.

28. Jedidi I, Zhang F, Qiu H, Stahl SJ, Palmer I, Kaufman JD, Nadaud PS, Mukherjee S, Wingfield PT, Jaroniec CP, Hinnebusch AG. 2010. Activator Gcn4 Employs Multiple Segments of Med15/Gal11, Including the KIX Domain, to Recruit Mediator to Target Genes in Vivo. J Biol Chem 285:2438–2455.

29. Swanson MJ, Qiu H, Sumibcay L, Krueger A, Kim S-J, Natarajan K, Yoon S, Hinnebusch AG. 2003. A multiplicity of coactivators is required by Gcn4p at individual promoters in vivo. Mol Cell Biol 23:2800–2820.

30. Yoon S, Qiu H, Swanson MJ, Hinnebusch AG. 2003. Recruitment of SWI/SNF by Gcn4p does not require Snf2p or Gcn5p but depends strongly on SWI/SNF integrity, SRB mediator, and SAGA. Mol Cell Biol 23:8829–8845.

31. Di Lello P, Jenkins LMM, Jones TN, Nguyen BD, Hara T, Yamaguchi H, Dikeakos JD, Appella E, Legault P, Omichinski JG. 2006. Structure of the Tfb1/p53 complex: Insights into the interaction between the p62/Tfb1 subunit of TFIIH and the activation domain of p53. Mol Cell 22:731–740.

32. Langlois C, Mas C, Di Lello P, Jenkins LMM, Legault P, Omichinski JG. 2008. NMR Structure of the Complex between the Tfb1 Subunit of TFIIH and the Activation Domain of VP16: Structural Similarities between VP16 and p53. J Am Chem Soc 130:10596–10604.

33. Knutson BA, Luo J, Ranish J, Hahn S. 2014. Architecture of the Saccharomyces cerevisiae RNA polymerase I Core Factor complex. Nat Struct Mol Biol.

34. Donczew R, Hahn S. 2017. Mechanistic differences in transcription initiation at TATA-less and TATA-containing promoters. Mol Cell Biol MCB. 00448–17.

35. Kuras L, Thomas D. 1995. Functional analysis of Met4, a yeast transcriptional activator responsive to S-adenosylmethionine. Mol Cell Biol 15:208–216.

36. Schwank S, Ebbert R, Rautenstrauss K, Schweizer E, Schüller HJ. 1995. Yeast transcriptional activator INO2 interacts as an Ino2p/Ino4p basic helix-loop-helix heteromeric complex with the inositol/choline-responsive element necessary for expression of phospholipid biosynthetic genes in Saccharomyces cerevisiae. Nucleic Acids Res 23:230–237.

37. Kolaczkowska A, Kolaczkowski M, Delahodde A, Goffeau A. 2002. Functional dissection of Pdr1p, a regulator of multidrug resistance in Saccharomyces cerevisiae. Mol Genet Genomics 267:96–106.

38. Stebbins JL, Triezenberg SJ. 2004. Identification, mutational analysis, and coactivator requirements of two distinct transcriptional activation domains of the Saccharomyces cerevisiae Hap4 protein. Eukaryotic Cell 3:339–347.

39. Ma J, Ptashne M. 1987. Deletion analysis of GAL4 defines two transcriptional activating segments. Cell 48:847–853.

40. Leuther KK, Salmeron JM, Johnston SA. 1993. Genetic evidence that an activation domain of GAL4 does not require acidity and may form a beta sheet. Cell 72:575–585.

41. Rothermel BA, Thornton JL, Butow RA. 1997. Rtg3p, a basic helix-loop-helix/leucine zipper protein that functions in mitochondrial-induced changes in gene expression, contains independent activation domains. J Biol Chem 272:19801–19807.

42. Thomas D, Jacquemin I, Surdin-Kerjan Y. 1992. MET4, a leucine zipper protein, and centromere-binding factor 1 are both required for transcriptional activation of sulfur metabolism in Saccharomyces cerevisiae. Mol Cell Biol 12:1719–1727.

43. Lee TA, Jorgensen P, Bognar AL, Peyraud C, Thomas D, Tyers M. 2010. Dissection of combinatorial control by the Met4 transcriptional complex. Molecular Biology of the Cell 21: 456–469.

44. Kuras L, Cherest H, Surdin-Kerjan Y, Thomas D. 1996. A heteromeric complex containing the centromere binding factor 1 and two basic leucine zipper factors, Met4 and Met28, mediates the transcription activation of yeast sulfur metabolism. EMBO J 15:2519–2529.

45. Blaiseau PL, Thomas D. 1998. Multiple transcriptional activation complexes tether the yeast activator Met4 to DNA. EMBO J 17:6327–6336.

46. Flick K, Ouni I, Wohlschlegel JA, Capati C, McDonald WH, Yates JR, Kaiser P. 2004. Proteolysis-independent regulation of the transcription factor Met4 by a single Lys 48-linked ubiquitin chain. Nat Cell Biol 6:634–641.

47. Flick K, Raasi S, Zhang H, Yen JL, Kaiser P. 2006. A ubiquitin-interacting motif protects polyubiquitinated Met4 from degradation by the 26S proteasome. Nat Cell Biol 8:509–515.

48. Ouni I, Flick K, Kaiser P. 2010. A transcriptional activator is part of an SCF ubiquitin ligase to control degradation of its cofactors. Mol Cell 40:954–964.

49. Tyrrell A, Flick K, Kleiger G, Zhang H, Deshaies RJ, Kaiser P. 2010. Physiologically relevant and portable tandem ubiquitin-binding domain stabilizes polyubiquitylated proteins. Proc Natl Acad Sci USA 107:19796–19801.

50. Ambroziak J, Henry SA. 1994. INO2 and INO4 gene products, positive regulators of phospholipid biosynthesis in Saccharomyces cerevisiae, form a complex that binds to the INO1 promoter. J Biol Chem 269:15344–15349.

51. Dietz M, Heyken W-T, Hoppen J, Geburtig S, Schüller H-J. 2003. TFIIB and subunits of the SAGA complex are involved in transcriptional activation of phospholipid biosynthetic genes by the regulatory protein Ino2 in the yeast Saccharomyces cerevisiae. Mol Microbiol 48:1119–1130.

52. Heyken W-T, Repenning A, Kumme J, Schüller H-J. 2005. Constitutive expression of yeast phospholipid biosynthetic genes by variants of Ino2 activator defective for interaction with Opi1 repressor. Mol Microbiol 56:696–707.

53. Kumme J, Dietz M, Wagner C, Schüller H-J. 2008. Dimerization of yeast transcription factors Ino2 and Ino4 is regulated by precursors of phospholipid biosynthesis mediated by Opi1 repressor. Curr Genet 54:35–45.

54. Klein P, Pawson T, Tyers M. 2003. Mathematical modeling suggests cooperative interactions between a disordered polyvalent ligand and a single receptor site. Current Biology.

55. Olsen JG, Teilum K, Kragelund BB. 2017. Behaviour of intrinsically disordered proteins in protein-protein complexes with an emphasis on fuzziness. Cell Mol Life Sci 12:269–9.

56. Flock T, Weatheritt RJ, Latysheva NS, Babu MM. 2014. Controlling entropy to tune the functions of intrinsically disordered regions. Curr Opin Struct Biol 26:62–72.

57. Sievers F, Wilm A, Dineen D, Gibson TJ, Karplus K, Li W, Lopez R, McWilliam H, Remmert M, Söding J, Thompson JD, Higgins DG. 2011. Fast, scalable generation of high-quality protein multiple sequence alignments using Clustal Omega. Molecular Systems Biology 7:539–539.

58. Alva V, Nam S-Z, Söding J, Lupas AN. 2016. The MPI bioinformatics Toolkit as an integrative platform for advanced protein sequence and structure analysis. Nucleic Acids Res 44:W410–5.

59. Sikorski RS, Hieter P. 1989. A system of shuttle vectors and yeast host strains designed for efficient manipulation of DNA in Saccharomyces cerevisiae. Genetics 122:19–27.

